# Revealing cell populations catching the early stages of the human embryo development in naïve pluripotent stem cells

**DOI:** 10.1101/2022.03.06.482812

**Authors:** Marta Moya-Jódar, Asier Ullate-Agote, Paula Barlabé, Juan Roberto Rodríguez-Madoz, Gloria Abizanda, Carolina Barreda, Xonia Carvajal-Vergara, Amaia Vilas-Zornoza, Juan Pablo Romero, Leire Garate, Xabier Agirre, Giulia Coppiello, Felipe Prósper, Xabier L. Aranguren

**Affiliations:** Program of Regenerative Medicine, Center for Applied Medical Research (CIMA), University of Navarra, Pamplona, 31008, Spain; Instituto de Investigación Sanitaria de Navarra (IdiSNA), Pamplona, 31008, Spain; Advanced Genomics Laboratory, Program of Hemato-Oncology, Center for Applied Medical Research (CIMA), University of Navarra, Pamplona, Spain; Hemato-Oncology Program, Center for Applied Medical Research (CIMA), IDISNA, University of Navarra, Pamplona, Spain; 10x Genomics, 6230 Stoneridge Mall Road, Pleasanton, CA 94588, USA; Hematology Department, Clínica Universidad de Navarra, University of Navarra, Pamplona, Spain; Centro de Investigación Biomédica en Red de Cáncer (CIBERONC), Pamplona, Spain

**Author notes:** co-first authors (Marta Moya-Jódar, Asier Ullate-Agote).

**Keywords:** Single-cell RNA-seq, Naïve state, Trophectoderm, 8CLCs

## Abstract

Naïve human pluripotent stem cells (hPSCs) are defined as the *in vitro* counterpart of the human preimplantation embryo’s epiblast and are used as a model system to study developmental processes. In this study, we report the discovery and characterization of distinct cell populations coexisting with epiblast-like cells in 5iLAF naïve human induced PSCs (hiPSCs) cultures. Noteworthily these populations closely resemble different cell types of the human embryo at early developmental stages. While epiblast-like cells represented the main cell population, interestingly we detected a cell population with gene and transposable element expression profile closely resembling the totipotent 8-Cell (8C) stage human embryo, and three cell populations analogous to trophectoderm (TE) cells at different stages of their maturation process: transition, early and mature stage. Thus, 5iLAF naïve hiPSCs cultures provide an excellent opportunity to model the earliest events of human embryogenesis, from the 8C stage to the peri-implantation period.

**Graphical abstract:** 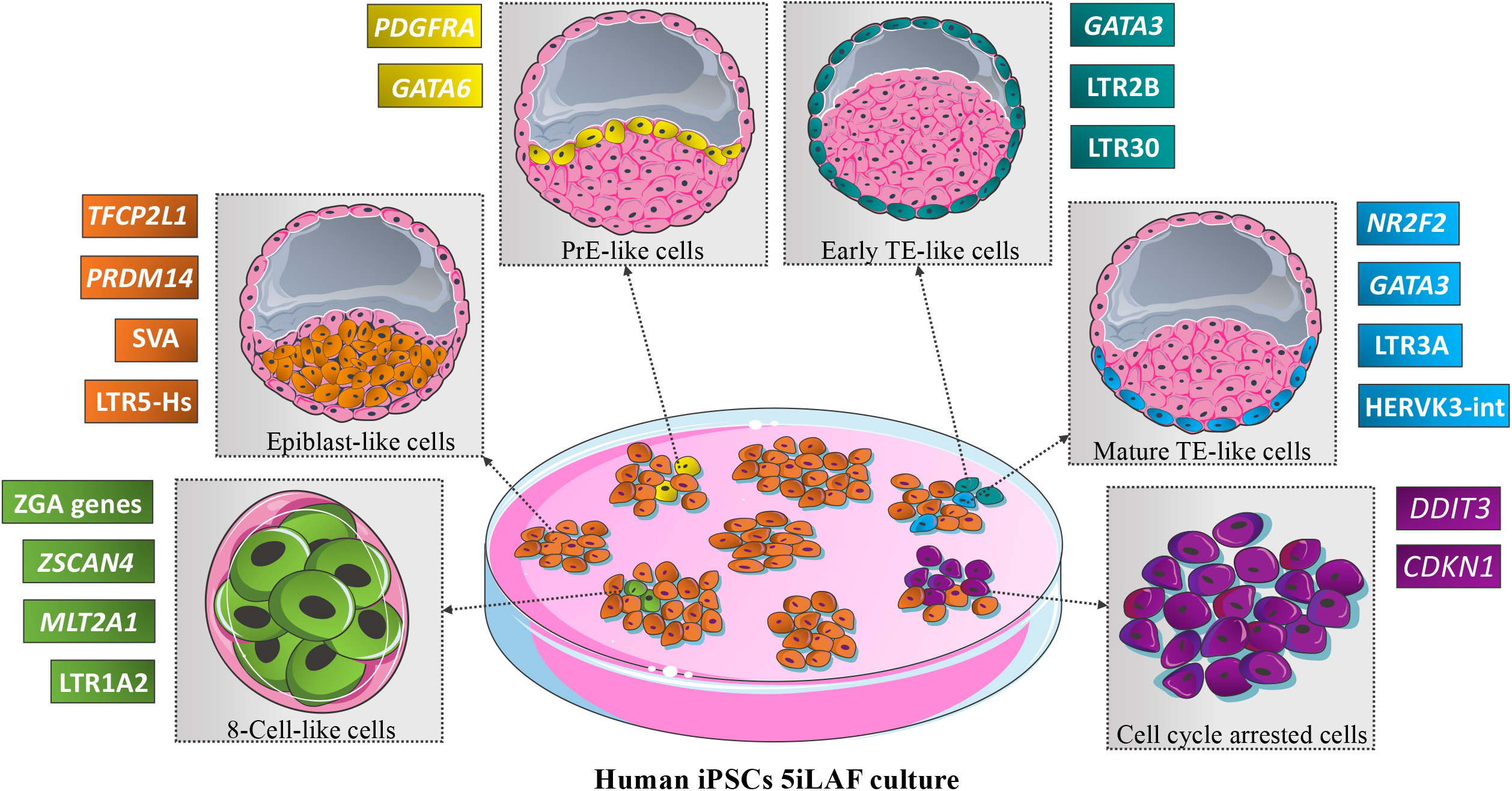

## INTRODUCTION

Embryonic development starts from a single totipotent cell called zygote, derived from the fertilized egg. Maternally derived RNA is used in the first phases of embryonic development, in particular, until the 2-Cell (2C) stage in mouse and the 8-Cell (8C) stage in human, when zygotic genome activation (ZGA) occurs, and the first wave of embryo genome transcription starts. In human, the first lineage differentiation events take place in the compact morula at day 4, with the outer cells initiating a trophectoderm-(TE) specific transcriptional program. As a result, 5 days after fertilization, the blastocyst is segregated into the inner cell mass (ICM), which will give rise to the embryo proper, and the TE, which supports uterine implantation. By day 6, the blastocyst’s ICM differentiates into the primitive endoderm (PrE), which will be a major constituent of the yolk sac, and the epiblast, which will form the fetus, while the TE will generate the extraembryonic tissues, including the placenta. Therefore, at the late blastocyst stage, prior to the implantation, these three main cell lineages are already determined (Shahbazi, 2020).

Human pluripotent stem cells (hPSCs) can be obtained and maintained either in primed state, representing the *in vitro* counterpart of the post-implantation epiblast, or in naïve state, corresponding to the pre-implantation epiblast. Diverse protocols have been established to obtain hPSCs with naïve characteristics (Chan et al., 2013; Gafni et al., 2013; Guo et al., 2016, 2017; X. Liu et al., 2017; Takashima et al., 2014; Theunissen et al., 2014; Ware et al., 2014). Comparative transcriptional and molecular analyses have demonstrated that, among these protocols, the 5iLA, t2i/L+Gö and PXGL culture conditions produce cells with the highest resemblance to the *in vivo* pre-implantation epiblast (Pastor et al., 2016; Stirparo et al., 2018; Takashima et al., 2014; Theunissen et al., 2014, 2016).

Certain studies have identified different subpopulations of naïve hPSCs in 5iLAF cultures based on the expression levels of SSEA4(Pastor et al., 2016); moreover, expression of the primitive endoderm (PrE) master gene GATA6 has been detected in different naïve cultures (Guo et al., 2016; Linneberg-Agerholm et al., 2019) and trophectoderm markers such as GATA3 and CDX2 have been identified in 5iLA cultures (Dong et al., 2020), suggesting the presence of different cell populations representing distinct cell lineages of the human pre-implantation embryo. However, a single cell transcriptomics study did not show significant heterogeneity in human embryonic stem cells (hESCs) in t2i/L+Gö naïve conditions (Messmer et al., 2019). Nevertheless, they interestingly identified a very small intermediate population presenting a gene expression profile that separate from both naïve and primed states (Messmer et al., 2019). On the other hand, there are significant evidences proving the existence of heterogeneous cell populations in mouse naïve PSCs cultures: cells equivalent to ICM (identified by the double expression of PDGFRα and CD31), PrE-precursors (PDGFRα^+^; CD31^-^)(lo Nigro et al., 2017) and 2-cell-like cells (2CLCs) (Macfarlan et al., 2012) have been described. 2CLCs are the *in vitro* equivalent to the 2C stage mouse embryo and are characterized by the expression of genes from the *Zscan4* family and MERVL retrotransposons, while showing downregulation of proteins associated with pluripotency, like Pou5f1, Sox2 and Nanog, a genetic signature that correlates with the ZGA taking place in the 2-cell mouse embryo (Macfarlan et al., 2012). In addition, a cell population transitioning from naïve to 2CLCs state was identified, being characterized by *Zscan4* genes expression, and lack of retrotransposon activation (Rodriguez-Terrones et al., 2018). Therefore, to fill the void of knowledge regarding hPSCs heterogeneity, in the present study we analyzed by single-cell RNA sequencing naïve human induced PSCs (hiPSCs) under 5iLAF culture conditions, finding different cell populations which capture distinct stages of human embryo development, from the 8C stage to the peri-implantation stage.

## RESULTS

### Identification of cell heterogeneity in naïve hiPSC cultures

To determine the cell heterogeneity of our 5iLAF naïve culture we performed single-cell RNA sequencing (scRNA-seq) and unsupervised clustering analysis of 3,652 cells, obtaining seven clusters (Figure 1A). First, we validated the naïve identity of our cells by integrating our data with previously published results (X. Liu et al., 2020), where the process of cell reprogramming from fibroblasts to iPSCs in primed and in naïve (RSeT and t2i/L+Gö) conditions was analyzed at single-cell resolution. Our sample clustered next to the naïve cells of this study and presented high score for naïve and epiblast gene signatures (Figure S1). These results were confirmed by morphology (Figure S2A), protein expression (Figure S2B-D), and a profound genome hypomethylation in LINE1 CpG sites (Figure S2E). Consistently, primed markers were not expressed in any of the seven clusters identified in our culture, demonstrating the lack of residual primed cells, except for CD24, which was enriched in Cluster 5 (Figure 1B). Naïve and pluripotency markers associated with the human embryo epiblast, were homogeneously expressed in Clusters 0 to 3, with the exception of *PRDM14*, that was lower in Cluster 2 (Figure 1C-D). On the other hand, some of these markers were downregulated in Clusters 4, 5 and 6 (Figure 1C-D). Additionally, correlation analysis of differentially expressed genes (DEG), showed high similarity between Clusters 0 to 3, while Clusters 4, 5 and 6 defined distinct cell populations (Figure 1E). Therefore, we considered Clusters 0 to 3, representing 80.5% of the total cells, as a single population which we named Epi-Cluster 0-3. A comparison of the DEG from this cluster (Table S1) with publicly available stage-specific gene expression modules of human embryos from zygote to late blastocyst stage (Stirparo et al., 2018), based on data from three different single-cell datasets20–22 showed that the majority of DEG (53.5%) were specific to either late (39.3%), or early (14.2%) blastocyst’s ICM (Figure 1F). In addition, we analyzed the expression of transposable elements (Table S2), as their transcription is regulated in a stage-specific manner during human early embryogenesis (Göke et al., 2015). In our Epi-Cluster 0-3 we observed high expression of SVA family members, as well as LTR5-Hs and HERVK-int elements (Figure 1G), which are characteristic of morula and early blastocyst stage, in line with previously described naïve cells-specific transposable elements (Theunissen et al., 2016).

**Figure 1:**
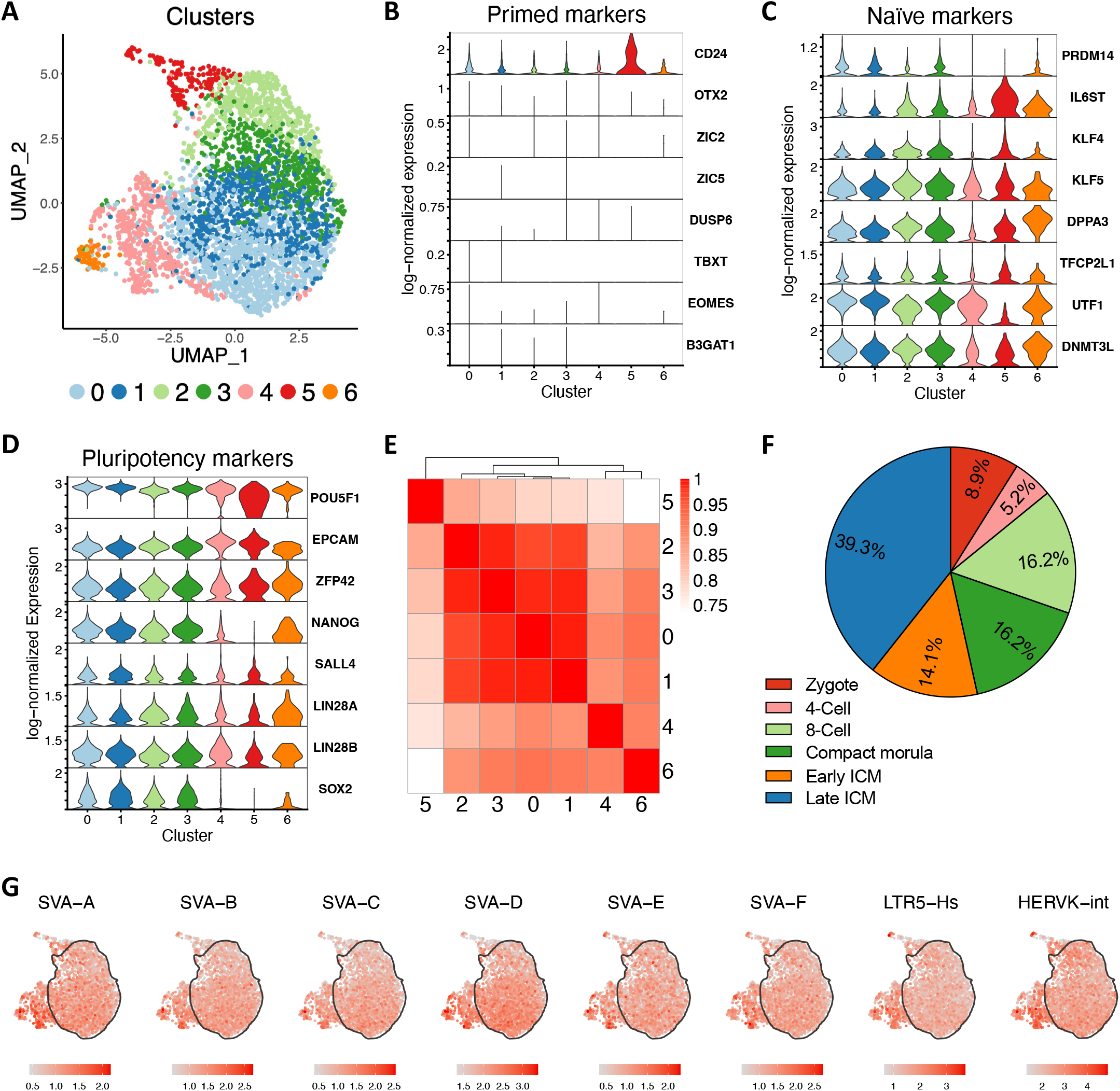
Overall analysis of cell heterogeneity in human naïve 5iLAF culture. (A) Uniform manifold approximation and projection (UMAP) of the 5iLAF-cultured hiPSCs (3,654 cells). Cells are color-coded according to unsupervised clustering analysis. (B-D) Violin plots showing single cell log-normalized gene expression of selected primed, naive and pluripotency markers in each cluster. Color code as in panel A. (E) Heatmap representing the correlation between the different clusters, considering differentially expressed genes in at least one cluster against the rest. (F) Proportion of differentially expressed genes in Epi-Cluster 0-3 corresponding to each stage-specific gene expression module of the human embryo at early developmental stages obtained from Stirparo et al., 2018 (Stirparo et al., 2018) and listed in their Table S6. Genes differentially expressed in Epi-Cluster 0-3 that were not specific to any developmental stage or were not annotated were remove from the analysis. (G) UMAP representations of the normalized expression of naïve-specific transposable elements. A contour delineates Epi-Cluster 0-3.

### Presence of trophectoderm-like cells in naïve hiPSC cultures

We found that cells in Cluster 5, which represent the 4.4% of total cells, displayed the highest scores for a previously described TE signature (Petropoulos et al., 2016) (Figure 2A). In agreement, known TE-associated genes (Petropoulos et al., 2016; Xiang et al., 2020) were enriched in this cluster (Figure 2B). Henceforth, we will refer to Cluster 5 as TE-like Cluster. RNA velocity analysis indicated that TE-like cells arose from naïve cells of Epi-Cluster 0-3 (Figure 2C). This finding is consistent with the described capacity of human naïve cells to differentiate into the TE lineage under defined culture conditions14,19,24–26. The presence of TE-like cells in our naïve culture was confirmed by the detection of GATA3^+^ cells by immunofluorescence staining (Figure 2D). Interestingly, within the TE-like Cluster, we were able to observe a few scattered cells expressing primitive endoderm (PrE)-specific genes (Stirparo et al., 2018): *PDGFRA, COL4A1, GATA6, GATA4* and *SOX17* (data not shown). The possible existence of PrE-like cells in naïve 5iLAF cultures is in line with the previously reported expression of GATA6 in human t2iLGÖY naïve cultures (Guo et al., 2016, 2017; Linneberg-Agerholm et al., 2019). Furthermore, PrE-like cells have been described in mouse ES cultures (lo Nigro et al., 2017).

**Figure 2:**
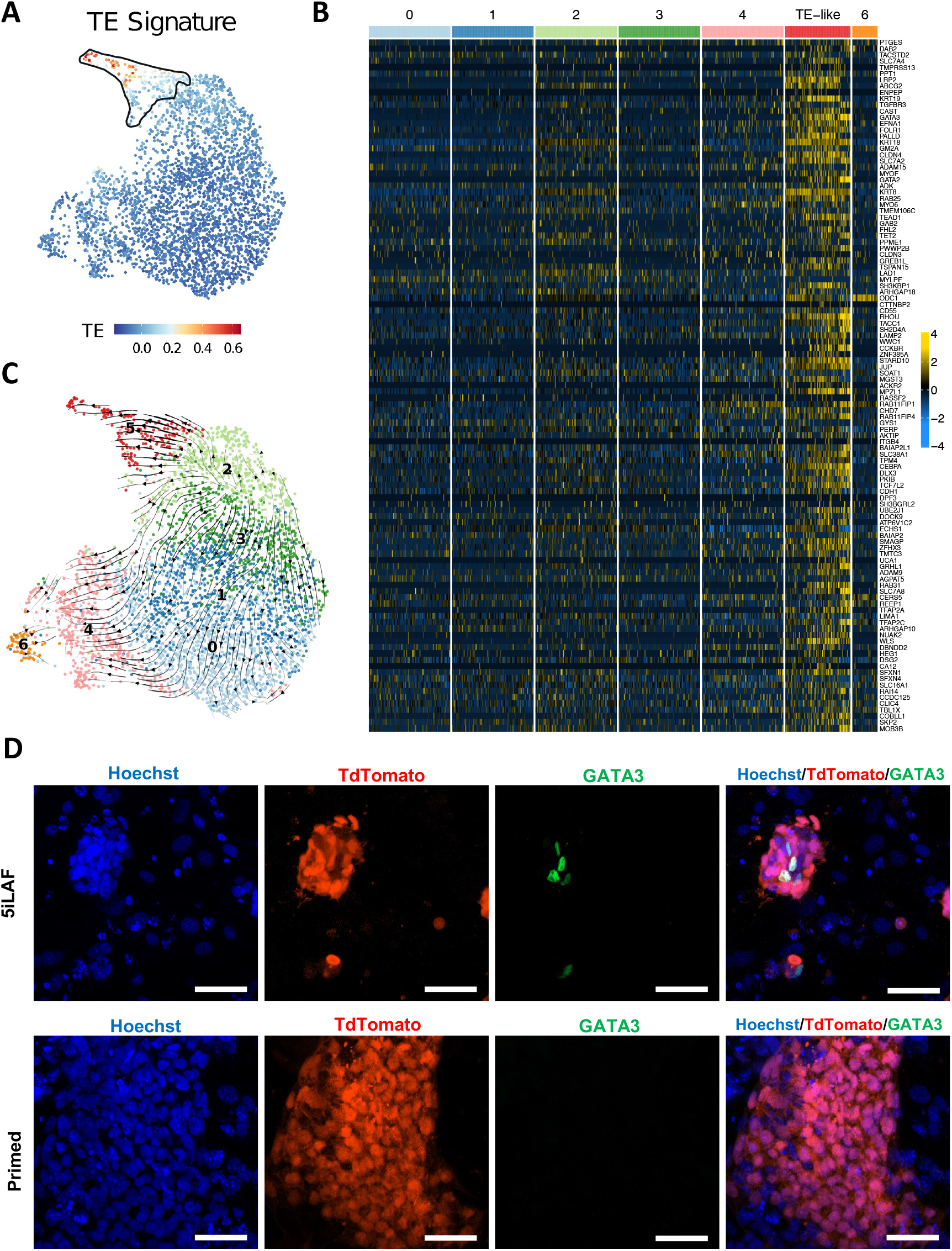
TE-like cells in human naïve 5iLAF culture. (A) UMAP representation of scores from a TE signature including TE-specific markers that are common between Petropoulos (Petropoulos et al., 2016) and Xiang (Xiang et al., 2020) studies. A contour delineates Cluster 5. (B) Heatmap representing scaled gene expression of the 111 genes from the TE signature. To facilitate visualization of the data, 200 cells from each cluster were randomly selected. Cells are ordered by cluster membership. (C) UMAP plot showing RNA velocities as streamlines. (D) Immunofluorescence staining of GATA3 (in green) in human naïve (upper panels) and human primed (lower panels) cultures. Human cells are labelled with tdTomato (in red) and Hoechst was used as a nuclear counterstain. Scale bars, 50 μm.

### hiPSCs naïve culture as a model of early TE specification process

Analysis of defined TE-associated markers showed heterogeneous expression within cells of the TE-like Cluster (Figures 2A and 3A). Accordingly, an unsupervised subclustering analysis revealed three subpopulations (Figure 3B). To determine possible differences in TE maturation stage between them, we scored the cells against the GATA2 and NR2F2 genes modules, recently described for early developmental stages of the human embryo (Meistermann et al., 2021). According to this study early TE cells (B3 stage) are GATA2-module^+^ and NR2F2-module^-^, while mature TE cells (B4-B6 stage) are GATA2-module^+^ and NR2F2-module^+^ (Meistermann et al., 2021). When applying the GATA2 module to our 5iLAF sample, we found the highest scores throughout the TE-like Cluster (Figure 3C). However, cells of Cluster 5_0 had relatively lower scores for this module, and mainly had negative scores for the NR2F2 module (Figure 3C), a first proof of their early maturation stage. Additionally, this cluster expressed epiblast markers such as *DPPA3, DPPA5, DNMT3L* and *ALPG* at higher level compared to the other two subclusters (Table S1). Co-expression of epiblast and early TE genes is a hallmark of TE transition in human embryonic development (Niakan et al., 2013; Stirparo et al., 2018), and thus we termed Cluster 5_0 as Transition-TE-like Cluster. Cluster 5_1 and Cluster 5_2 could be distinguished by differences in scores for the NR2F2-module, being higher in the latter cluster (Figure 3C). Therefore, we named Cluster 5_1 as Early-TE-like Cluster and Cluster 5_2 as Mature-TE-like Cluster. To corroborate these results, we applied signatures derived *in vitro* for the different steps of the differentiation process of TE cells from pre-cytotrophoblasts stage (pre-CTBs, corresponding to developmental day 6-7 of the human embryo; D6-7) to extravillous trophoblasts (EVTs; D14)(Xiang et al., 2020). Cells with the highest scores for the pre-CTBs (D6-7) signature were found in the Transition- and Early-TE-like Clusters, while cells in the Mature-TE-like Cluster presented the highest scores for the Early syncytiotrophoblast (STB; D8-10) signature (Figure 3D). On the other hand, cells from later stages of placental differentiation were not present, according to the low scores obtained for their corresponding signature (data not shown) and the absence of placental polypeptide hormones expression (Figure S3A). Therefore, these results confirm that the three subclusters found in the TE-like Cluster define TE-like cells at different maturation stages and thus naïve hiPSCs in 5iLAF medium can be a model to study the transition from epiblast cells to early and mature TE cells. For instance, the modulation of the transcription factors (TFs) differentially expressed in our TE-like subclusters (Figure S3B-C and Table S1) could elucidate their role in the TE-differentiation process. Moreover, in our 5iLAF cultures we retrieved 38 transposable elements differentially expressed in the TE-like Cluster, when compared to the rest of cells. Between them, we identified repetitive sequences from the LINE1, Alu and LTR families (Figure 3E-F and Table S2). Noteworthily 86 transposable elements were differentially expressed in Early- or Mature-TE-like subclusters compared to the other clusters, suggesting a specific role of those sequences in the maturation process of TE cells. Between the previously described TE-specific transposable elements for example, we found that LTR30, LTR10A (L. Liu et al., 2019) and LTR2B, were enriched in the Early-TE-like Cluster. On the other hand, LTR3A (L. Liu et al., 2019) HERVK3-int and LTR10B2 were enriched in the Mature-TE-like Cluster (Table S2). Finally, transposable elements known to be involved in placental development (Senft et al., 2021) such as endogenous retroviruses ERVW-1 and ERVFRD-1 were enriched in the Mature TE-like Cluster (Table S1). Thus, the study of transposable elements could help to elucidate the process of TE-maturation.

**Figure 3:**
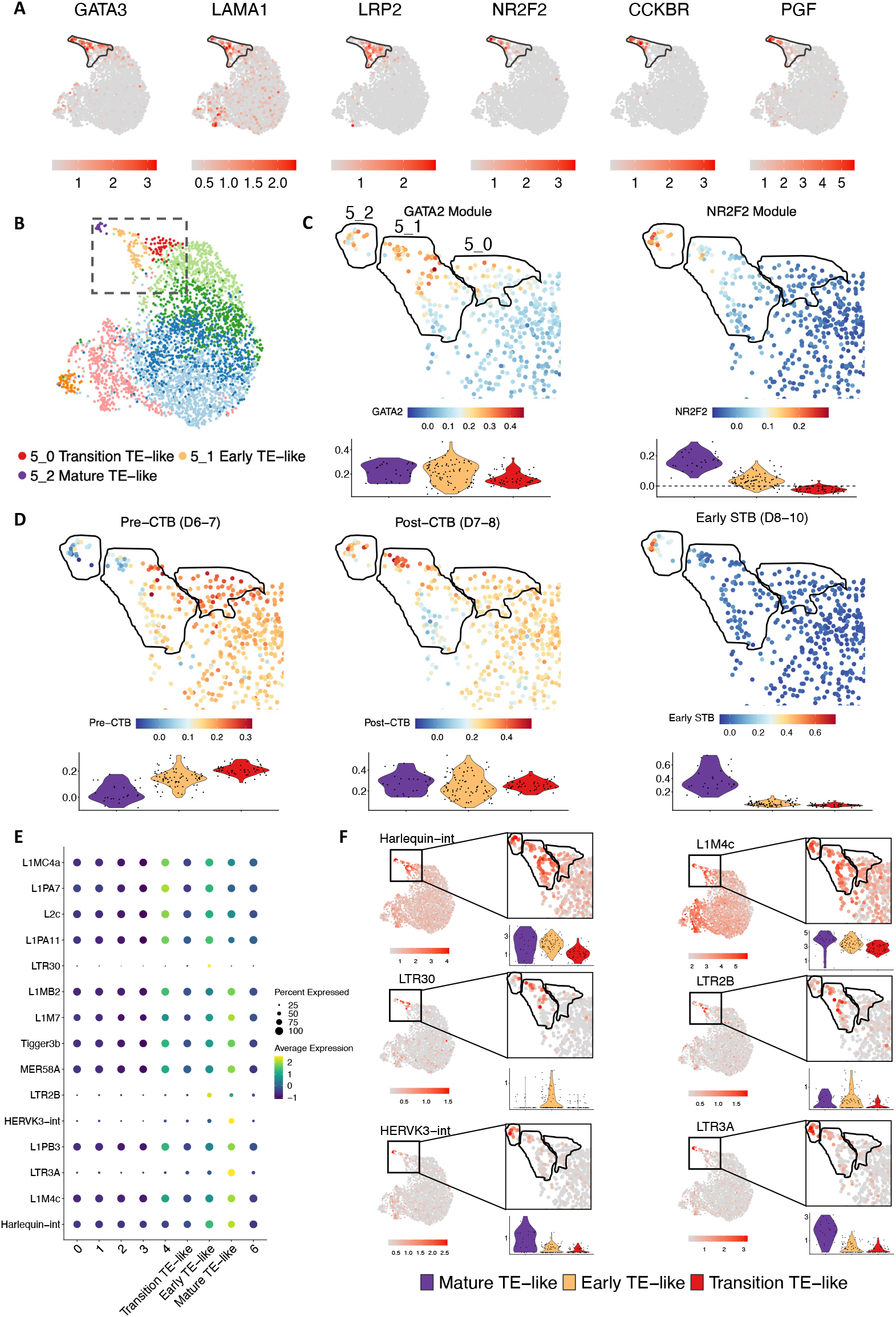
Analysis of the three different stages of TE maturation present in the TE-like Cluster. (A) UMAP representing log-normalized expression of a selected group of TE-associated markers. A contour delineates TE-like Cluster. (B) UMAP representation after subclustering the TE-like Cluster. Cells are color-coded according to unsupervised clustering analysis. The dashed box corresponds to the UMAP region shown in the upcoming panels C and D. (C) UMAP representation of scores from the GATA2 gene module and the NR2F2 gene module signatures from Meistermann (Meistermann et al., 2021). A contour delineates each subpopulation from the TE-like Cluster. Violin plots of the scores grouped by TE-like subpopulations are depicted underneath, with a dashed line at score zero for the NR2F2 module. (D) UMAP representation of gene signature scores from different trophoblast lineages obtained from Xiang (Xiang et al., 2020): pre-CTB (days 6-7), post-CTBs (days 7-8) and early STBs (days 8-10). A contour delineates each subpopulation from the TE-like Cluster. Violin plots of the scores grouped by TE-like subpopulations are depicted underneath. (E) Dot plot representing scaled average expression of the top differentially expressed transposable elements in each of the TE-like subclusters. The size of the dot represents the proportion of cells expressing such element in each cluster. (F) UMAP representing the log-normalized expression in TE-like subclusters of selected transposable elements. A zoom-in of the TE-like cluster is displayed, with a contour delineating each subpopulation. Violin plots grouped by TE-like subpopulations are shown underneath the zoom-ins. The color code from the legend is the same for the violin plots in panels C, D and F.

Maturation of TE cells is associated with specific metabolic requirements (Kaneko, 2016; Posfai et al., 2019), so next we analyzed the metabolic transcriptome in the different subclusters. Interestingly, pathways related to energy production were enriched in Early-TE-like Cluster (Figure S3D and Table S1), in agreement with the high ATP consumption of these cells *in vivo*, required to expand the blastocoel cavity (Houghton et al., 2003). Meanwhile, Mature-TE-like Cluster was enriched in pathways related to steroid hormone synthesis/metabolism and regulation of angiogenesis, all functions needed to allow proper implantation of the embryo into the uterine wall (Figure S3D and Table S1). Finally, molecules playing a key role in the embryo attachment to the endometrium (Idelevich et al., 2020) were observed in Mature-TE-like cells, such as *PGF, CGA, TGFB1, VEGFA* (Table S1).

In conclusion, TE-like cells present in naïve cultures mimic the early stages of TE development and could be an ideal model to study the transition from epiblast to TE, TE maturation, and the role of these cells in the implantation process *in vitro*.

### Identification of 8-cell like cells (8CLCs) in 5iLAF culture

Interestingly, in Cluster 6 of our analysis, which corresponds to 1.7% of the total cells, we found cells expressing *ZSCAN4* (Figure 4A), a defining gene of the 8C stage human embryo (Stirparo et al., 2018). Additionally, in this cluster we detected expression of *DUXA, PRAMEF1-2, LEUTX, KLF17*, and other markers associated with the human 8C stage undergoing the ZGA process (Jiang et al., 2002; Maeso et al., 2016; Stirparo et al., 2018; Töhönen et al., 2015; Wang et al., 2018) (Figure 4A and Table S1). On the other hand, we observed significantly lower expression levels of *NANOG, SOX2*, and *PRDM14* compared to the Epi-Cluster 0-3, but not *ZFP42* or *POU5F1* (Figure 1C-D). This is in accordance with previous studies describing the downregulation of pluripotency-associated markers such as *Nanog, Sox2, Prdm14* and *Tfap2c*, but not *Pou5f1* during the transition from ESCs to 2CLCs in mouse cultures (Rodriguez-Terrones et al., 2018). The expression of *ZFP42* together with *ZSCAN4* supports the notion that Cluster 6 does not emerge from a process related to cell differentiation, since this process requires loss of *ZFP42* expression (Rodriguez-Terrones et al., 2018). To further determine to which developmental stage of the human embryo Cluster 6 corresponds to, we compared its DEG to human embryo transcriptomic data from zygote to late blastocyst stage (Stirparo et al., 2018). From the annotated genes (Stirparo et al., 2018), we observed that ∼40% were associated with the 8C stage and ∼20% with compact morula (Figure 4B). To refine this comparison, we next applied to our sample the DUXA module, which consists of genes associated with the ZGA between the 8C and morula stage in humans (Meistermann et al., 2021). Consistently with our previous analysis we found the highest scores in Cluster 6 (Figure 4C). On the other hand, as DUX4 has been recently identified as a key regulator of the ZGA genetic signature both *in vivo* and *in vitro* (de Iaco et al., 2017), we also applied the DUX4-induced signature (Hendrickson et al., 2017) to our data, and again we observed that Cluster 6 was enriched for these genes (Figure 4D). These data consistently show that Cluster 6 gene expression profile closely resemble that of the 8C human embryo undergoing ZGA, therefore we termed it 8C-like Cluster.

**Figure 4:**
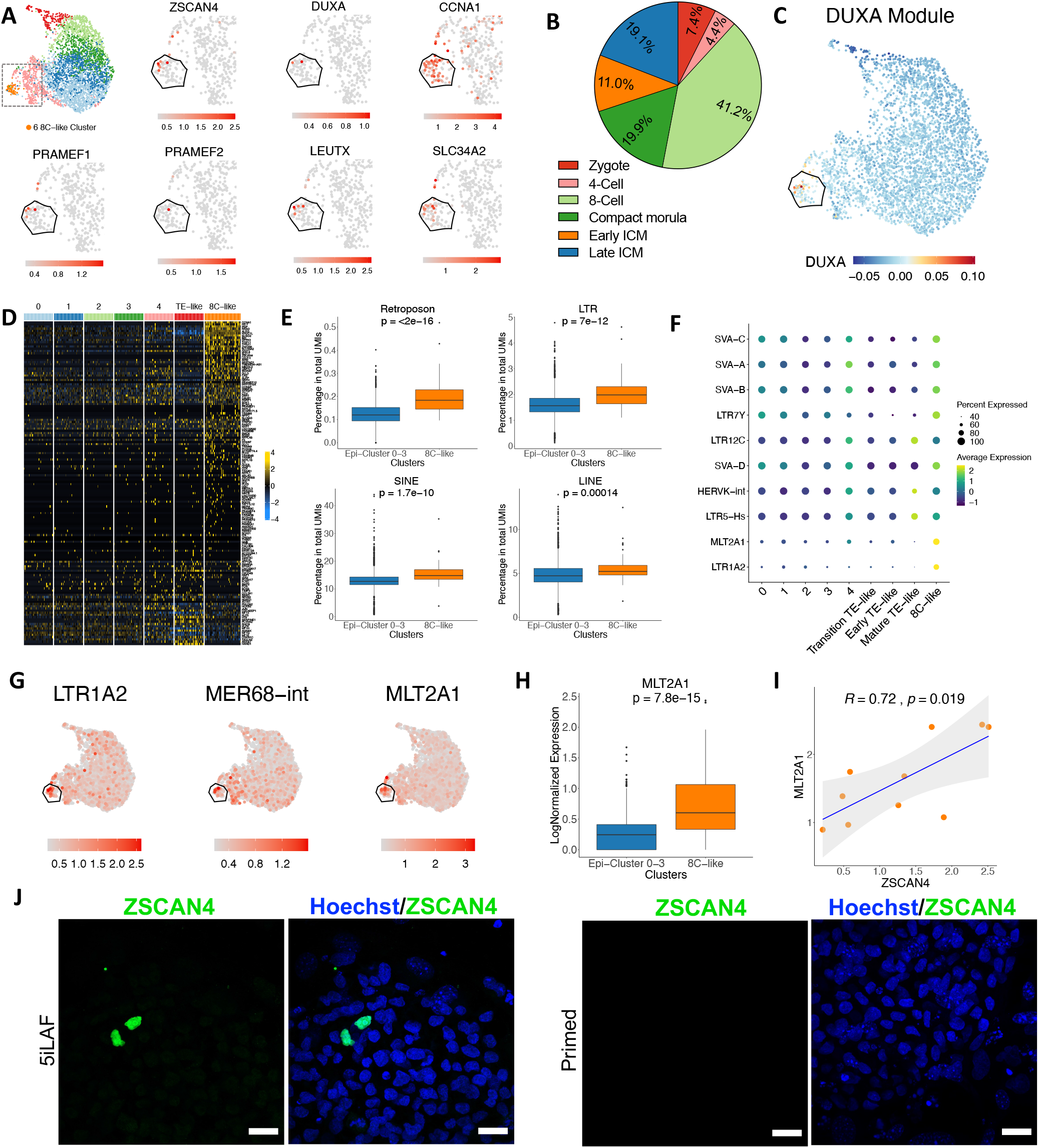
8CLCs in human naïve 5iLAF culture. (A) Zoom-ins of the UMAP region including the 8C-like Cluster representing the log-normalized expression of selected 8C embryo associated markers. The zoom-ins correspond to the dashed region shown in the top left UMAP. A contour delineates the 8C-like Cluster. (B) Proportion of differentially expressed genes in the 8C-like Cluster corresponding to each stage-specific gene expression module of the human embryo at early developmental stages obtained from Stirparo (Stirparo et al., 2018) and listed in their Table S6. Genes differentially expressed in 8C-like Cluster that were not specific to any developmental stage or were not annotated were removed from the analysis. (C) UMAP representation of scores from the DUXA gene module from Meistermann (Meistermann et al., 2021). A contour delineates the 8C-like Cluster. (D) Heatmap representing scaled gene expression of 144 markers upregulated after DUX4 induction in hiPSCs from Hendrickson (Hendrickson et al., 2017). All cells from 8C-like Cluster were represented, while to facilitate visualization of the data, 50 cells from each other cluster were randomly selected. Cells are ordered by cluster membership. (E) Boxplots showing the percentage of total UMIs corresponding to different transposable element classes (retroposons, LTRs, SINEs and LINEs) in cells from Epi-Cluster 0-3 and the 8C-like Cluster. (F) Dot plot representing scaled average expression of the top 10 differentially expressed transposable elements in the 8C-like Cluster. The size of the dot represents the proportion of cells expressing such element in each cluster. (G) UMAP representation of log-normalized expression of transposable elements that are enriched in the 8C-like Cluster. A contour delineates the 8C-like Cluster. (H) Boxplots showing the percentage of total UMIs corresponding to the MLT2A1 LTR element in cells from Epi-Cluster 0-3 and 8C-like Cluster. Two-side Wilcoxon Rank Sum tests were performed to obtain statistical significance in the comparisons between clusters from panels E and H (I) Correlation analysis between *ZSCAN4* and MLT2A1 expression in *ZSCAN4*^+^ cells from the 8C-like Cluster. (J) Confocal microscopy images of immunofluorescence staining for ZSCAN4 (in green) in 5iLAF (on the left) and primed (on the right) hiPSCs. DNA was counterstained with Hoechst. Scale bars, 20 μm.

As the expression of MERVL retrotransposons is a hallmark of 2CLCs in mouse (Macfarlan et al., 2012), we also analyzed transposable elements expression in the 8C-like Cluster. In this cluster we observed a significantly higher proportion of reads corresponding to retroposons, LTRs, SINEs and LINEs families compared to Epi-Cluster 0-3 (Figure 4E). Additionally, transposable elements known to be expressed between 8C and morula stages in human embryos such as the SVA retroposons, LTR5-Hs, and LTR7Y (Göke et al., 2015; L. Liu et al., 2019) were enriched in the 8C-like Cluster, as well as 8C-specific transposable elements such as HERVK-int, LTR12C and MLT2A1 (Göke et al., 2015; Grow et al., 2015; L. Liu et al., 2019). Moreover, we found that Mer68-int and LTR1A2, which have not been previously described as being expressed in early phases of embryonic development, were highly expressed in the 8C-like Cluster.

Noteworthily, MLT2A1, the most specific repetitive sequence to the 8C human embryo *in vivo* (Göke et al., 2015), was among the most enriched in the 8C-like Cluster (Fig 4F-H and Table S2). Importantly, *ZSCAN4* expression correlated with MLT2A1 expression (Figure 4I). Thus, we defined the *ZSCAN4*^*+*^*/*MLT2A1^*+*^cells found in the 8C-like Cluster as 8CLCs. In mPSC cultures, *Zscan4*^+^/MERVL^+^ 2CLCs range from 0.1 to 0.5% of the total cells (Macfarlan et al., 2012). Accordingly, in our naïve 5iLAF sample, we observed a similar proportion of 8CLCs (∼0.3%). Finally, we confirmed by immunofluorescence staining the existence of ZSCAN4^+^ cells in 5iLAF cultures, while they were absent in primed cultures (Figure 4J).

In conclusion, these data prove the existence of 8CLCs in 5iLAF cultures. These cells could be an ideal *in vitro* platform to study the earliest stages of the human embryo development, including the ZGA events.

### Presence of a cell population characterized by cell cycle arrest

Cluster 4, representing 13.4% of the total cells, expressed several epiblast markers, although could be distinguished by a very low expression of *NANOG, SOX2, PRDM14* and *KLF4* (Figure 1C-D). The most prominent characteristic found in this cluster was the high percentage of cells in cell cycle arrest (G1 phase, Figure 5A). In fact, the most significantly enriched gene sets for this cluster are related to response to DNA damage and cell cycle arrest (Figure 5B). When studying the differential expression of transposable elements in Cluster 4, we found LINE elements of the L1-subfamily, among others (Figure 5C-D). Interestingly, L1 elements play an important role in genomic instability and cancer and L1 retrotransposition induces cell stress via DNA response to damage(Grundy et al., 2021). The transcriptional profile of Cluster 4 suggests cell damage and is consistent with the known detrimental effects on genetic and epigenetic stability induced by naïve conditions on cultured hPSCs (X. Liu et al., 2017; Pastor et al., 2016; Theunissen et al., 2014).

**Figure 5:**
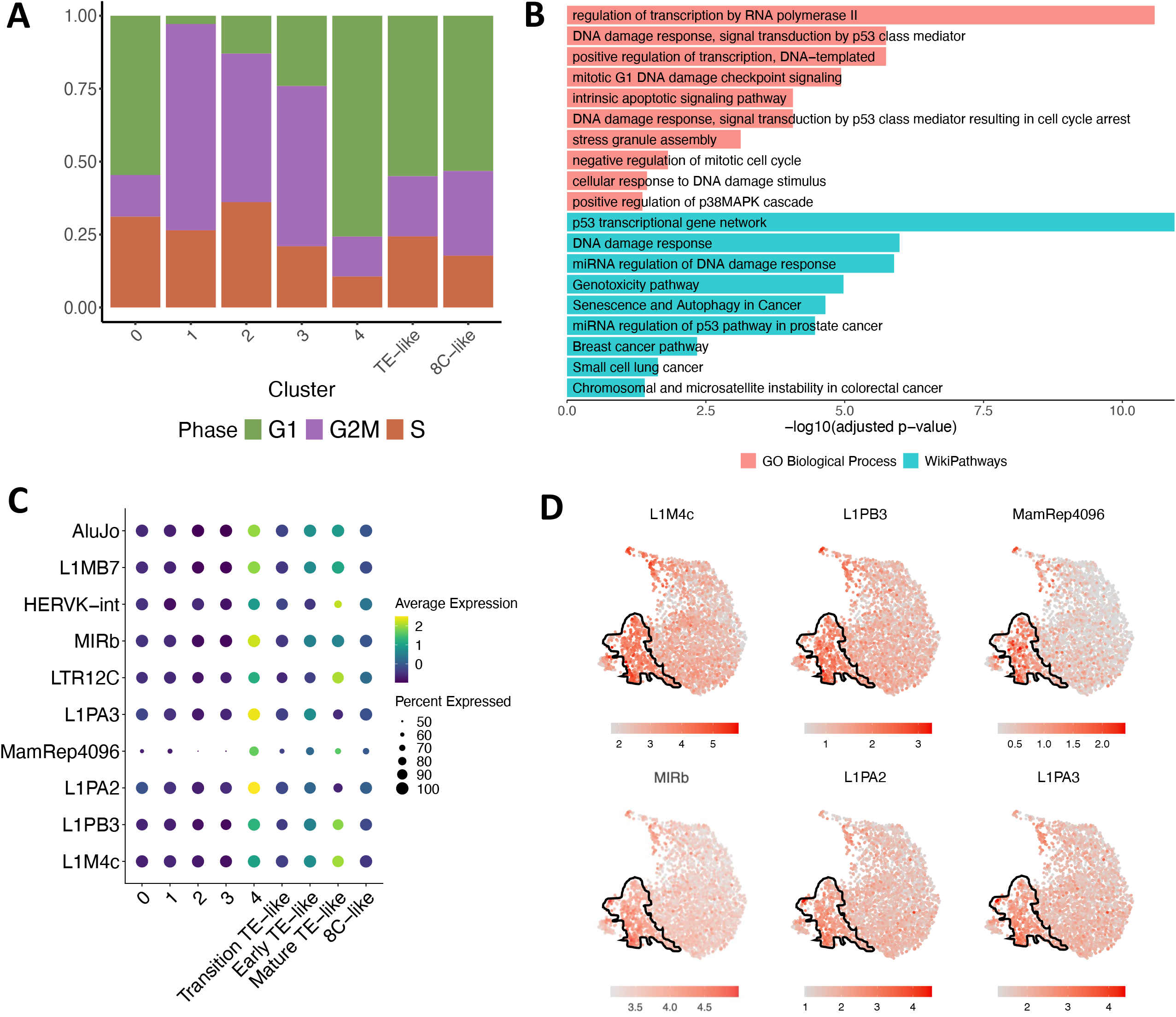
Presence of naïve cells with arrested cell cycle in Cluster 4. (A) Proportion of cells in each cell cycle phase for the unsupervised clusters. (B) Selection of significant GO Biological Processes and WikiPathways terms from an overrepresentation analysis using DEG in Cluster 4 against the rest. (C) Dot plot representing scaled average expression of the top 10 differentially expressed transposable elements in Cluster 4. The size of the dot represents the proportion of cells expressing such element in each cluster. (D) UMAP representation of log-normalized expression of differentially expressed transposable elements in Cluster 4, delineated by a contour.

### Heterogeneous contribution of hiPSCs to human-mouse chimera

Epiblast-like naïve PSCs microinjected in a host embryo are known to contribute to the ICM (Beddington et al., 1989). To functionally prove the existence of cells corresponding to distinct developmental stages in our cultures, we conducted chimera formation experiments. We microinjected 8-10 5iLAF cells, labelled with tdTomato, into morula-stage mouse embryos and cultured them for 48 h *in vitro* (Figure 6A-B). Quantification at 24 h showed that part of the microinjected cells did not engraft in the host embryo, as we observed a decrease in cell number at this time point, whereas we detected cell proliferation at 48 h, as indicated by an increased number of cells (Figure 6C). Strikingly, at this time point, in addition to finding human cells in the ICM, we also observed a fraction of cells contributing to the TE (Figure 6D-F). TE identity of the human cells located in the outer layer of the chimeric embryos was confirmed by immunofluorescence for the GATA3 marker (Fig. 6D-E). These results further support the existence of cell heterogeneity within human 5iLAF cultures.

**Figure 6.**
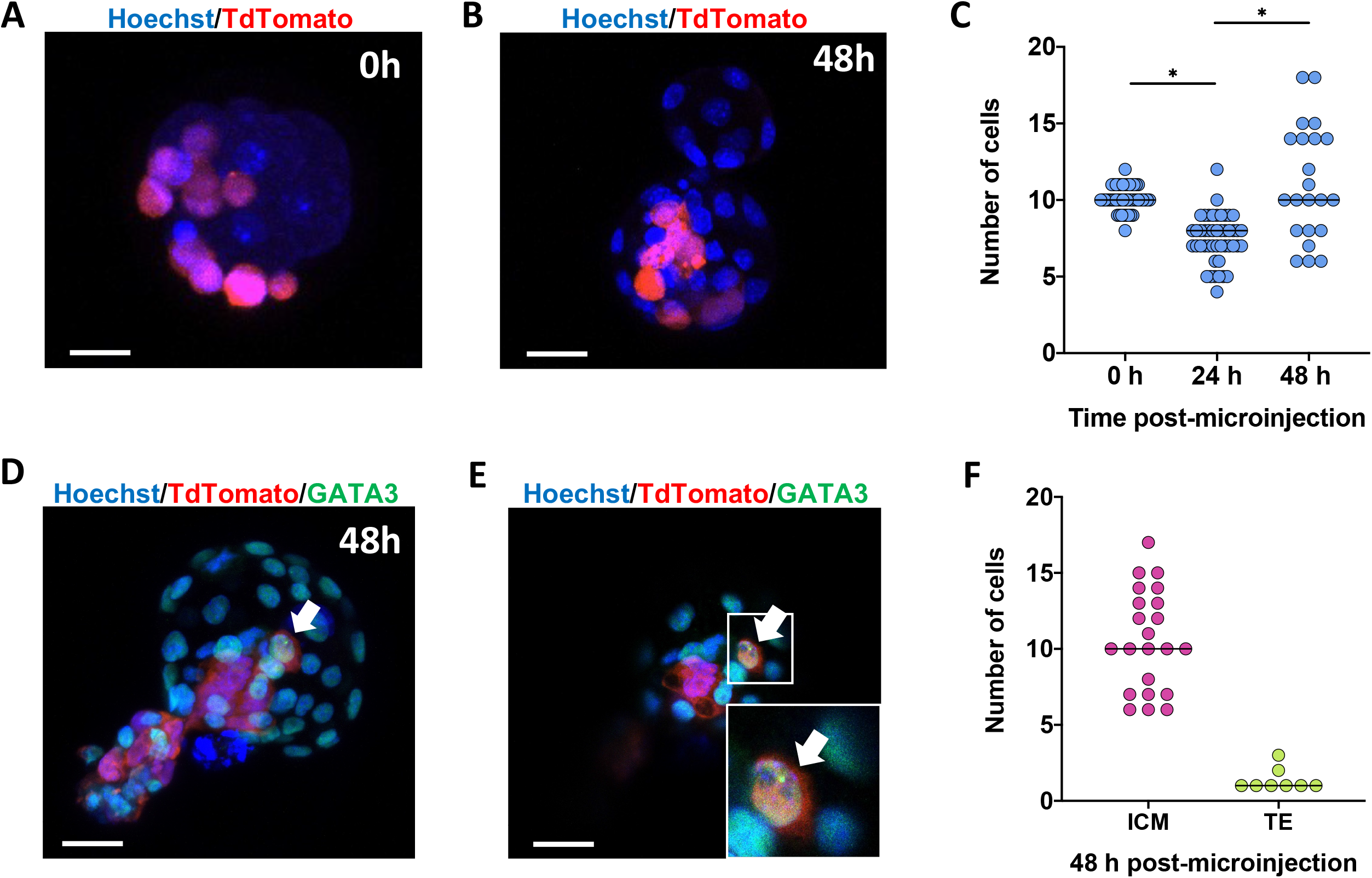
Contribution of human 5iLAF cells to the TE lineage in human-mouse chimera embryos. (A) Orthogonal projection of a mouse morula freshly microinjected with 10 human 5iLAF tdTomato^+^ cells. (B) Engraftment of human cells within the mouse embryo after 48 h of *in vitro* culture, shown by orthogonal projection confocal image. (C) Graphical representation of the human cells present into mouse embryos over time. (D) Immunostaining of GATA3 (in green) in a mouse embryo microinjected with human 5iLAF cells and cultured *in vitro* for 48 h. (E) Orthogonal section from panel D. Boxed area is enlarged in the bottom right corner of the picture. The arrow indicates a tdTomato^+^ human cell expressing GATA3 in panels D and E. (F) Graphical representation of human cells contribution to ICM or TE 48 h post-microinjection. Dots represent microinjected embryos. Hoechst was used as a nuclear counterstain in A, B, D and E. Scale bars, 20 μm.

## DISCUSSION

With this work we reveal and thoroughly describe four main cell populations with distinct characteristics coexisting in human naïve 5iLAF cultures. While the expression profile of 80% of the cells, here named Epi-Cluster 0-3, defined them as the *in vitro* counterpart of the late blastocyst epiblast cells, which corresponds to the gold standard definition of naïve, we also revealed the presence of TE-like cells and 8CLCs, as well as a population of Epiblast-like cells in cell cycle arrest.

The TE-like Cluster is characterized by a gene expression profile comparable to the early TE of the human embryo (D6-9) and expresses at high levels TE-specific transposable elements. This population represents 4.4 % of the cells. Such population of cells was not previously found in naïve PXGL conditions, possibly as a consequence of the presence of WNT and aPKC inhibitors, which block the access to TE differentiation in this medium(Guo et al., 2021). RNA velocity trajectory analysis showed that cells of the TE-like Cluster arise from naïve cells. This is in line with the known potential of naïve hPSCs to give rise to trophoblast stem cells (hTSCs) in a defined media (Okae et al., 2018), being hTSCs the *in vitro* counterpart of the trophoblast at D8-12 (Castel et al., 2020; Cinkornpumin et al., 2020; Dong et al., 2020; Guo et al., 2021; Io et al., 2021).

Interestingly, TE-like cells found in our naïve cultures presented a heterogeneous gene and transposable elements expression profile, that allowed us to subdivide them into three subclusters, corresponding to three early extraembryonic tissue maturation stages: transitional, early TE and mature TE. Thus, while hTSCs can be used to study trophoblast lineages differentiation (Castel et al., 2020; Cinkornpumin et al., 2020; Dong et al., 2020; Guo et al., 2021; Io et al., 2021), TE-like cells in 5iLAF cultures, which resemble the trophectoderm of the blastocyst at day D6-9, may be a suitable model to shape the role of the different genetic determinants of TE formation and its early maturation.

TE cells contribute to the process of implantation by modulating endometrial receptivity (Posfai et al., 2019). In the TE-like Cluster, we observed factors previously described to take part in this process and we unraveled new candidates that may play an important role in trophoblast to endometrium communication as, for instance *SEMA4C* (Table S1), whose receptor, PLEXIN B2, has been reported to participate in endometrial integrity (Singh et al., 2015). Since trophoblast dysfunction leads to complications during pregnancy, such as pre-eclampsia and intrauterine growth restriction, the study of TE-like cells present in patient-derived naïve iPSC cultures would facilitate the identification of the molecular players inducing these complications.

Human 5iLAF cells microinjected at the morula stage into mouse embryos showed contribution to both ICM and TE, with TE-contribution present in 36% of the chimeric embryos at 48 h post-injection. The identification of hiPSCs TE-contribution importantly confirms the existence of cells with *in vivo* TE-differentiation capacity in human naïve culture. We hypothesize that these cells are more likely to be derived from the TE-like Cluster, as the proportion of 8CLCs in our culture is too low to justify such a high frequency of TE-contribution. Accordingly, in chimera experiments with mouse PSCs, where a population of 2CLCs (Macfarlan et al., 2012) but not a TE-like population has been described, the contribution of microinjected cells to the TE was extremely rare (Beddington et al., 1989). Nevertheless, to prove this hypothesis, it would be necessary to identify surface markers that allow the isolation of TE-like cells from the naïve bulk and to perform chimera formation experiments with these isolated cells.

Noteworthily, 8C-like Cluster defines a population of cells enriched with the expression of 8C embryo specific genes and transposable elements, while showing downregulation of pluripotency-associated genes like *PRDM14, SOX2* and *NANOG*. In this population we detected the existence of *ZSCAN4*^*+*^/MLT2A1^+^ cells, which present a gene and transposable elements expression characteristics of the 8C stage of the human embryo *in vivo*. These 8CLCs represent ∼0.3% of the total cells, consistent with the described proportion of 2CLCs in mouse cultures (Macfarlan et al., 2012). This discovery is in line with the recent description of an analogous 8CLCs population in hESCs cultured naïve PXGL medium (Taubenschmid-Stowers et al., 2022). This observation is a further demonstration of the analogy of hiPSC with hESCs and opens the door to the use of naïve PSCs as a model to study key pathways determining the origin and developmental potential of 8CLCs. The *in vivo* counterpart of these cells is characterized by a higher state of pluripotency compared with ICM cells, therefore it would be of interest to test the chimeric capacity of 8CLCs with functional experiments. However, the isolation of this cell population and its expansion would be needed in order to perform these studies. For that, it will be of pivotal importance to identify suitable surface markers or generate reporter cell lines and to find key molecules to obtain an adapted culture media.

Unlike mouse PSCs, which are genetically stable after prolonged culture (Cervantes et al., 2002; Suda et al., 1987), human naïve PSCs are prone to acquire mutations, chromosomal abnormalities and epigenetic aberrations (Pastor et al., 2016; Theunissen et al., 2014) and therefore is still necessary to revise the naïve culture conditions to improve cell fitness. Accordingly, in our study we have observed that cells of Cluster 4 are in cell cycle arrest and express genes related to response to DNA damage. This observation reinforces the notion that renewed refinement of 5iLAF medium is required to improve cell stability. Stress response related genes enriched in Cluster 4, such as *DDIT3* and *CDKN1A* (Ock et al., 2020) (Table S1), could be used as markers to monitor cell integrity upon modification of the culture conditions.

In summary, our study describes the heterogeneity present in 5iLAF naïve hiPSCs. Distinct populations corresponding to 8CLCs and TE have been found to coexist with epiblast-like (naïve) cells in these cultures. Hence, 5iLAF condition could be used as a model to investigate human embryo development from the 8C stage, when the zygotic genome activation event occurs, to the peri-implantation period, where the correct maturation of TE cells play a key role, as well as to broaden the understanding of stem cell biology, thus representing an alternative to the use of human embryos and the associated ethical issues.

## ACKNOWLEDGMENTS

This study was supported by the Instituto de Salud Carlos III co-financed by European Regional Development Fund-FEDER “A way to make Europe” Red de Terapia Celular TERCEL (RD16/0011/0005) and Centro de Investigación Biomédica en Red de Cáncer CIBERONC (CB16/12/00489), and co-financed by European Union – NextGenerationEU. Redes de Investigación Cooperativa Orientada a Resultados en Salud RICORS (RD21/0017/0009 and RD21/0017/0019), the Ramón y Cajal State Program (MINECO/FSE, RYC-2015-17233), Ministerio de Ciencia e Innovación/Agencia Estatal de Investigación/FEDER, (UE/RTI2018-064485-B-I00), MINECO/FSE, (IJCI-2017-33070) and Gobierno de Navarra I+D 2020 Co-Funded by FEDER Funds (0011-1365-2020-000293 and 0011-1365-2020-000287).

## AUTHOR CONTRIBUTION

M.M.J. Generated and characterized naive hiPSCs, performed single-cell experiment, performed mouse-human chimera experiments, analyzed data, and wrote the manuscript. A.U. Performed bioinformatic analysis of the data, and wrote the manuscript. P.B. Provided assistance with experiments and edited the manuscript. A.V., G.A., C.B, and L.G. provided assistance with experiments. J.P.R. provided assistance with bionformatic analysis. J.R.R., X.C.V and X.Ag. provided materials and technical advice and edited the manuscript. G.C. discussed the data and wrote the manuscript. F.P. and X.A. conceived the research, provided funding, supervised the experiments, analyzed the data, and wrote the manuscript.

## AUTHOR DISCLOSURE STATEMENT

J.P.R is an employee and shareholder of 10x Genomics.

## STAR METHODS SECTION

### RESOURCE AVAILABILITY

#### Lead Contact

Further information and requests for resources and reagents should be directed to and will be fulfilled by the Lead Contact, Xabier L. Aranguren.

#### Animal models

All animals were housed in specific pathogen free conditions with free access to food and water. All animal procedures were performed according to all state and institutional laws, guidelines and regulations. All studies were approved by the Ethics Committee for Animal Research at the University of Navarra and the Government of Navarra.

#### Cell culture

G15.AO human-induced pluripotent stem cell (hiPSC) line was used in this study (Zapata-Linares et al., 2016). Cells were routinely maintained on irradiated mouse embryonic fibroblast (iMEF) feeders in primed conditions: DMEM Knock-Out (Gibco #10829-018), 20% Knock-Out Serum Replacement (KSR, Gibco #10828-028), 1% GlutaMAX (Gibco #35050-038), 1% non-essential amino acids (Gibco #11140-035), 1% Antibiotic-Antimycotic (Gibco #15240-062), 0.1 mM β-mercaptoethanol (Gibco #31350-010) and 20 ng/mL human FGF-basic (Peprotech #100-18B). After reaching 70% confluence (5-7 days), cells were passaged by detaching them with 200 U/mL collagenase type IV (Gibco #17104-019) for 10 minutes, followed by mechanical disruption of the colonies with cell scrapper and gentle pipetting. Cells were cultured in a humidified atmosphere at 37ºC, 20% O_2_, 5% CO_2_ in air.

#### 5iLAF medium

HiPSCs were converted into naïve-like cells as described (Theunissen et al., 2014) with minor modifications of the media: briefly, 1:1 Neurobasal (Gibco #21103-049), DMEM-F12 (Sigma-Aldrich #D6421), 1X B27 Supplement (Gibco #17504-044), 1X N2 Supplement (Gibco #17502-048), 1% GlutaMAX (Gibco #35050-038), 1% Non-Essential Amino Acids (Gibco #11140-035), 1% Antibiotic-Antimycotic (Gibco #15240-062) 0.5% Knock-Out Serum Replacement (KSR Gibco #10828-028), 20 ng/mL recombinant human LIF (Peprotech #300-05-100UG), 20 ng/mL recombinant human Activin A (Peprotech #120-14-10UG), 8 ng/mL human FGF-basic (Peprotech #100-18b-50UG), and the following small molecule inhibitors: 10 μM Y27632 (Axon Medchem #1683); 1 μM WH-4-23 (Biovision #2827-10), 1 μM PD0325901 (Axon Medchem #1408), 1 μM IM-12 (Sigma-Aldrich #SML0084-5MG); 0.5 μM SB590885 (TOCRIS #2650). Briefly, hiPSCs were passaged every 7–9 days by single-cell disaggregation with 1:1 TrypLE (Gibco #12605-028) 1X PBS/0.25 mM EDTA (Invitrogen #AM9260G) and plated on an iMEF feeder layer. Culture medium was supplemented with additional Y27632 inhibitor for the first 24 h at 5 μM. Cells were cultured in a humidified atmosphere at 37ºC, 5% O_2_, 5% CO_2_ in air and checked monthly for Mycoplasma contamination.

#### Cells immunofluorescence staining

Primed and naïve hiPSCs were cultured over iMEFs in 8-well chambers (Thermo Scientific Nunc#177445) until they reached 60-70% of confluence. Then, they were fixed with 4% paraformaldehyde (PFA) solution in PBS for 30 minutes at 4°C, washed 3 times in PBS, permeabilized with PBS 0.1% Triton™ X-100 (Fisher BioReagents #BP151-500) for 1 h at 4°C, washed again and then, blocked in PBS 1% BSA (Sigma-Aldrich # A3912-100G) for 1 hour at 4°C. Samples were stained with the following primary antibodies in blocking solution overnight at 4°C: Anti-POU5F1 (1:50, Santa Cruz Biotechnology #sc-9081), Anti-SOX2 (1:50, Sigma-Aldrich #AB5603), Anti-TFE3 (1:200, Merck -Sigma Aldrich #HPA023881-100UL), Anti-hTFCP2L1 (1:200, R&D systems #AF5726), Anti-GATA3 (1:100, Abcam #ab199428), Anti-NANOG (1:50, ab80392, abcam), and Anti-ZSCAN4 (1:100, Origene #TA800535). After rinsing three times with PBS to remove the unbound antibodies, samples were exposed to the corresponding Alexa Fluor™ 488 goat anti-rabbit IgG secondary antibody (Invitrogen #A11008), Alexa Fluor® 488 donkey anti-goat IgG secondary antibody (Life Technologies #A11055) or Alexa Fluor® 488 goat anti-mouse IgG secondary antibody (Invitrogen #A11029), for 2 hours at 4°C in darkness (dilution 1:500 in blocking solution). After 3 washing steps with PBS, nuclear counterstain was performed with Hoechst 33342 trihydrochloride (Invitrogen #H21492) at 1:800 dilution for 30 minutes at 4°C in darkness. Images were recorded on a Confocal Scanning Laser Microscope (Zeiss LSM 800) and ZEN 2.3 system software. Composite images were obtained with ZEN 2.3 system software.

#### Flow Cytometry and FACS sorting

Pluripotency-, primed-, and naïve-associated markers were analyzed by FACS. Briefly, hiPSCs were cultured up to 70% confluence and washed once with PBS before single-cell dissociation. Then, hiPSCs were dissociated by incubation with TrypLE-EDTA (1:1 TrypLE-0.25mM mM EDTA/PBS) for 5 min. HiPSCs were resuspended in 100 μL of 1% BSA-PBS and incubated with the corresponding antibody for 30 min at 4 °C in the dark: BV510-conjugated Mouse Anti-human CD24 at 1:100 dilution (BD Horizon/ BD Biosciences #563035), BV421-conjugated Mouse Anti-human CD57 at 1:100 dilution (BD Horizon/ BD Biosciences #563896), PE-conjugated anti-human CD90 at 1:100 dilution (BD Biosciences #555596), APC-conjugated Mouse Anti-human SSEA4 at 1:100 dilution (R&D Systems #FAB1435A), BV510-conjugated Mouse Anti-human CD77 at 1:100 dilution (BD Horizon/ BD Biosciences #563630), APC-conjugated Mouse Anti-human CD130 at 1:40 dilution (BioLegend #362006). Isotype-matched IgGs were used as negative controls. Thereafter, samples were washed twice with PBS and finally resuspended in FACS buffer (PBS 1mM EDTA, 25 mM HEPES, 1% BSA, pH7), to be recorded in a BD FACSCanto™ II with BD FACSDiva software. Flow cytometry data were analyzed using FlowJo software.

#### Lentiviral production and cell transduction

For lentiviral particle production, HEK293 cells were plated 5×10^6^ cells/10 cm dish (Corning #430167) and the next day transfected with the FUtdTW TdTomato reporter plasmid (Addgene #22478) together with two helper plasmids (psPax2 and PMD2G) using X-tremeGENE HP DNA Transfection Reagent (Roche). In brief, 400 μL of OPTI-MEM (Invitrogen) was mixed with 1 μg PMD2G, 3 μg psPax2 and 4 μg lentiviral construction plasmid. 24 μL of X-tremeGENE HP DNA Transfection Reagent was added and the mixture was incubated for 20 minutes at room temperature and gently applied to the cells. The next day, medium was replaced and lentiviral particle-containing supernatant was collected 36 hours later. Lentiviral particles were concentrated by ultracentrifugation (2 h at 20.000 g at 4°C) and resuspended in PBS. Cells were transduced with the lentiviral particles, and 7 days later, tdTomato positive (tdTomato^+^) cells were sorted using a BD FACSAria™ Ilu, analyzed with BD FACSdiva software.

#### Mouse morula microinjection with 5iLAF hiPSCs

Fertilized mouse embryos were harvested from superovulated B6.DBA2 female mice crossed with males of the same strain at 2.5 days postcoitum (dpc) by flushing the infundibulum with M2 medium (Sigma-Aldrich #M7167-100ML). Human cells were dissociated with TrypLE-PBS-EDTA and magnetically selected to eliminate dead cells (Miltenyi Biotech #130-090-101) and remove mouse feeders (Miltenyi Biotech #130-095-531) through LS Columns (Miltenyi Biotech #130-042-401) following the manufacturer’s instructions. Ten tdTomato^+^ 5iLAF hiPSCs were microinjected into the perivitelline space of the morula using Leica DMI3000 B microscope and mechanical micromanipulators with hanging joystick; TransferMan® NK2 (Eppendorf). After microinjection, embryos were cultured in 5iLAF medium for 48 h at 37°C under 5% O_2_, 5% CO_2_ in air (Drawer Type Incubator #AD-3100), until they developed to the blastocyst stage. Then, tdTomato^+^ hiPSCs inside morulas and blastocysts were photographed and quantified by eye under a fluorescence microscope. We statistically analyzed our sample using Kruskal-Wallis test with an adjusted p-value < 0.05.

#### Mouse embryos immunofluorescence staining

Mouse embryos 0-, 24- and 48-h post-microinjection with human naïve 5iLAF cells were fixed with 4% PFA solution in PBS for 45 min at room temperature, washed in PBS and permeabilized with PBS 0.1% TritonX-100 for 2 h at 4 °C. Then, embryos were blocked in PBS 1% BSA (Sigma-Aldrich # A3912-100G) for 1 h at 4 °C. Samples were stained with GATA3 (Abcam #ab199428) at 1:50 in blocking solution overnight at 4°C. After rinsing with PBS to remove the unbound antibodies, samples were exposed to the secondary antibody; Alexa Fluor™ 488 goat anti-rabbit IgG (Invitrogen #A11008) for 2 hours at 4°C in darkness (dilution 1:500 in PBS). Nuclear counterstain was performed with Hoechst 33342 trihydrochloride (1:800) for 30 min at 4°C in darkness. Images were recorded with Confocal Scanning Laser Microscope (Zeiss LSM 800) and ZEN 2.3 system software. The contribution of 5iLAF hiPSCs to the trophectoderm linage was determined by co-expression of tdTomato and GATA3 within mouse embryos’ orthogonal sections. Composite images were obtained with ZEN 2.3 system software.

#### Pyrosequencing

Pyrosequencing assay was designed to analyze DNA methylation in five CpG dinucleotides of LINE-1 human retrotransposon in primed and naïve hiPSCs. Sodium bisulfite modification of 1 μg of genomic DNA was carried out with the CpGenome™ DNA Modification Kit (Chemicon #S7820) following the manufacturer’s protocol. After bisulfite treatment of DNA, PCR was performed using PyroMark master mix (QIAGEN) with a denaturalization at 95°C for 15 min and for 45 cycles consisting of denaturation at 94°C for 1 min, annealing at 55°C for 1 min, and extension at 72°C for 1 min, followed by a final 10 min extension. The PCR reactions for pyrosequencing were done with a biotinylated specific reverse primer (LINE-Forward: 5’-TTTTGAGTTAGG TGTGGGATATA-3’and LINE-Reverse: 5’-Biotin-AAAATCAAAAAATTCCCTTTC-3’). The resulting biotinylated PCR products were immobilized to streptavidin Sepharose High Performance beads (GE healthcare, Uppsala, Sweden) and processed to yield high quality ssDNA using the PyroMark Vacuum Prep Workstation (Biotage, Uppsala, Sweden), according to the manufacturer’s instructions. The pyrosequencing reactions were performed using the sequencing primer (LINE-Seq: 5’-AGTTAGGTGTGGGA TATAGT-3’) in the PyromarkTM ID (QIAGEN). Sequence analysis was performed using the PyroQ-CpG analysis software (QIAGEN). Human male genomic DNA universally methylated for all genes (Intergen Company, Purchase, NY) was used as a positive control for methylated alleles. Water blanks were included with each assay.

#### Single-cell RNA-sequencing (scRNA-seq)

Twenty-four hours before single-cell dissociation, 5iLAF human cells were treated with rock inhibitor Y27632 (5 μM - Axon Medchem #1683). Next day, human cells were dissociated using 1:1 TrypLE (Gibco #12605-028) 1X PBS and 0.25 mM EDTA (Invitrogen #AM9260G) for 5 min at 37 °C and resuspended in FACS buffer (1X PBS, 1 mM EDTA, 25 mM HEPES pH=7 and 1% BSA). Then, tdTomato^+^ human cells were FACS sorted with BD FACSAria™ IIu through a nozzle of 100 μm in 1X PBS 0.05% BSA, and the number of cells was quantified in a Neubauer chamber.

The transcriptome of tdTomato+ naïve hiPSCs cultured under 5iLAF conditions for 3 passages was examined using NEXTGEM Single Cell 3’ Reagent Kits v3 (10x Genomics) according to the manufacturer’s instructions. 15,000 cells were loaded at a concentration of 700-1000 cells/μL on a Chromium Controller instrument (10X Genomics) to generate single-cell gel bead-in-emulsions (GEMs). In this step, each cell was encapsulated with primers containing a fixed Illumina Read 1 sequence, a cell-identifying 16 bp 10x barcode, a 10 bp Unique Molecular Identifier (UMI) and a poly-dT sequence. Upon cell lysis, reverse transcription yielded full-length, barcoded cDNA. This cDNA was then released from the GEMs, PCR-amplified and purified with magnetic beads (SPRIselect, Beckman Coulter). Enzymatic Fragmentation and Size Selection was used to optimize cDNA size prior to library construction. Fragmented cDNA was then end-repaired, A-tailed and ligated to Illumina adaptors. A final PCR-amplification with barcoded primers allowed sample indexing. Library quality control and quantification was performed using Qubit 3.0 Fluorometer (Life Technologies) and Agilent’s 4200 TapeStation System (Agilent), respectively. Sequencing was performed in a NextSeq500 (Illumina) (Read1: 28 cycles; Read2: 49 cycles; i7 index: 8 cycles) at an average depth of 45,000 reads/cell. The sequenced library was demultiplexed and converted to FASTQ files using the mkfastq function from Cell Ranger v5.0.1 (10x Genomics). Reads were aligned to the reference human genome (GRCh38) downloaded from the 10x Genomics website (version 2020-A) using the Cell Ranger (v5.0.1) count function with default parameters. Genome annotation corresponded to Ensembl v98. The median number of unique molecular identifiers (UMIs) and genes detected per cell were 20,302 and 4,537 respectively.

#### scRNA-Seq analysis

The computational analysis of the resulting UMI count matrices was performed using the R package Seurat (v 4.0.5) (Stuart et al., 2019). Cells were subjected to a quality control step, keeping those cells expressing between 1,000 and 6,500 genes, that had between 2,000 and 48,000 UMIs and with less than 10% of UMIs assigned to mitochondrial genes. Those thresholds were chosen upon visual inspection of distributions. A preliminary clustering retrieved a fibroblast population that was also removed from further analysis. Using this filtering, we kept 3,652 cells, with a median of 4,851 genes per cell. Genes detected in less than 3 cells were removed from the analyses.

The dataset was subjected to normalization, identification of highly variable features and scaling using the SCTransform function (Hafemeister et al., 2019), regressing out for the % of mitochondrial UMIs. To characterize the cell populations present, we performed an unsupervised clustering analysis using the Louvain algorithm with a resolution of 0.5 in a shared nearest neighbors graph constructed with the first 20 principal components, as implemented in the FindClusters and FindNeighbors Seurat functions. Non-linear dimensional reduction for visualization was done using the RunUMAP function with the same principal components. Cluster 5 was further subdivided using the FindSubCluster function with a 0.3 resolution. Cluster markers were identified using the FindAllMarkers function in the log-normalized counts by using the Wilcoxon Rank Sum test, keeping as differentially expressed those that presented an average log2 fold-change greater than 0.25 and an adjusted p-value < 0.05. To perform overrepresentation analyses the R package enrichR (Kuleshov et al., 2016) was used with several databases, including GO_Biological_Process_2021 and WikiPathways_Human_2021. Signatures scores were obtained using the AddModuleScores function from Seurat. Genes in each signature were converted to their corresponding symbol from Ensembl v98. Cell cycle analysis was performed using the CellCycleScoring function with default parameters. The plots were generated using the DimPlot, DotPlot, VlnPlot and FeaturePlot functions from Seurat as well as the ggplot2, ComplexHeatmap and pheatmap R libraries.

#### Integrated analysis with Liu dataset

The scRNASeq datasets from Liu et al.(X. Liu et al., 2020) (RSeT, naïve, primed and days 0-7 of reprogramming) and this study were combined. Genes expressed in less than 10 cells and with an average expression lower than 0.01 were discarded. UMI counts were normalized using the NormalizedData function in Seurat and principal component analysis was performed using 2,000 variable features. The Harmony algorithm (Korsunsky et al., 2019) was used to integrate the samples with default parameters as implemented in the RunHarmony function, using the library correspondence as the only batch information for correction. UMAP embeddings were obtained using the first 30 batch-corrected principal components.

#### RNA velocity

The Cell Ranger output was processed using velocyto v0.17.17 (la Manno et al., 2018) with the ‘run10x’ mode to obtain a loom file including a count matrix of spliced and unspliced read counts. We merged this matrix together with a loom file generated from the Seurat analysis as input for scVelo v0.2.4(Bergen et al., n.d.) choosing there the top 2,000 variable genes that share a minimum of 30 counts for spliced and unspliced mRNA. We learned the transcriptional dynamics of splicing kinetics and estimated the RNA velocity with the ‘recover_dynamics’ and ‘velocity’ (in dynamical mode) functions. We embedded the resulting velocities on the UMAP space obtained with Seurat with the ‘velocity_embedding_stream’ function.

#### Transposable element analysis

We used scTE (He et al., 2021) to quantify the expression of transposable elements. First, we built an index with the repetitive elements from hg38 in bed format and the annotation used for Cell Ranger (Ensembl v98) with scTE_build. Then, we ran scTE with the index and the output BAM file from Cell Ranger, obtaining a counts matrix including both genes and repetitive elements. We processed the matrix with Seurat as above, keeping the same cells and assigning the same clusters to find differentially expressed transposable elements and proportions of different families of repetitive elements.

## SUPPLEMENTAL INFORMATION

**Supplementary Figure 1. Integrated analysis with scRNASeq data for human PSC reprogramming under naïve and primed conditions**. (A) UMAP representation after integrating the single cell data from this study (in green), with the scRNASeq libraries from Liu et al. (X. Liu et al., 2020), comprising data from fibroblasts (D07) and human PSC reprogramming process under primed and naïve (t2i/L+Gö and RSet) conditions. Plot is color-coded by library correspondence. (B-D) UMAP representation of the scores from primed (X. Liu et al., 2020), epiblast(Petropoulos et al., 2016) and naïve(X. Liu et al., 2020) gene signatures in the integrated dataset. (E-H) UMAP representations of the log-normalized expression of fibroblast (E), primed (F), pluripotency (G) and naïve associated (H) markers in the integrated dataset. Contour delineates cells from our sample in all panels.

**Supplementary Figure 2. 5iLAF cells characterization**. (A) Phase contrast images of primed and 5iLAF naïve hiPSCs. Scale bars, 200 μm. (B-C) Confocal microscopy images of tdTomato-labeled primed and 5iLAF naïve hiPSCs immunostained for (B) the pluripotency markers SOX2 and POU5F1 and (C) the naïve markers TFCP2L1 and TFE3. DNA was counterstained with Hoechst. Scale bars, 50 μm. (D) Histograms of flow cytometry analysis for primed- (CD24, CD57, CD90 and SEEA4) and naïve-(CD77 and CD130) state surface markers on hiPSCs cultured in primed and 5iLAF naïve mediums. (E) DNA methylation levels of specific LINE1 CpG sites in primed and 5iLAF naïve hiPSCs.

**Supplementary Figure 3. Analysis of the different TE-like subclusters**. (A) Heatmap representing scaled gene expression of a set of polypeptide hormones expressed in different trophoblast lineages from Xiang et al. (Xiang et al., 2020). Cells are ordered by subcluster membership. (B) Heatmap representing scaled gene expression of differentially expressed markers when comparing a TE-like subcluster against the other two. Cells are ordered by subcluster membership. On the right, list of differentially expressed transcription factors. (C) UMAP representations of the log-normalized expression of transcription factors expressed in TE-like subclusters. A zoom-in of the TE-like Cluster is displayed, with a contour delineating each subpopulation. Violin plots grouped by TE-like subpopulations are shown underneath the zoom-ins. (D) Selection of significant GO Biological Processes and WikiPathways terms from an overrepresentation analysis using DEG for the Early TE-like Cluster and the Mature TE-like Cluster when compared against each other.

