## Supplementary figures and images for "Revealing cell populations catching the early stages of the human embryo development in naïve pluripotent stem cells"

### Supplementary Figures

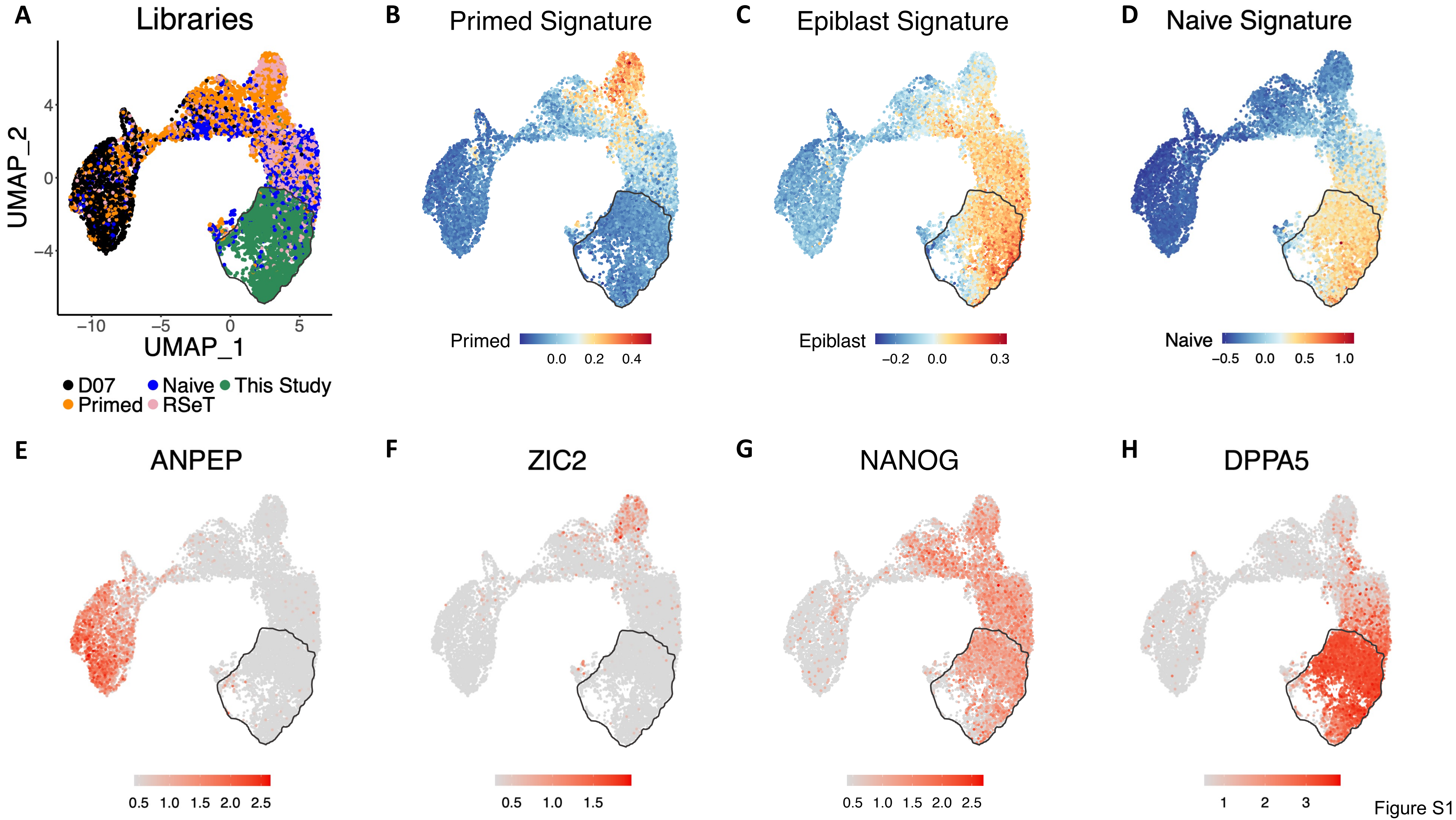

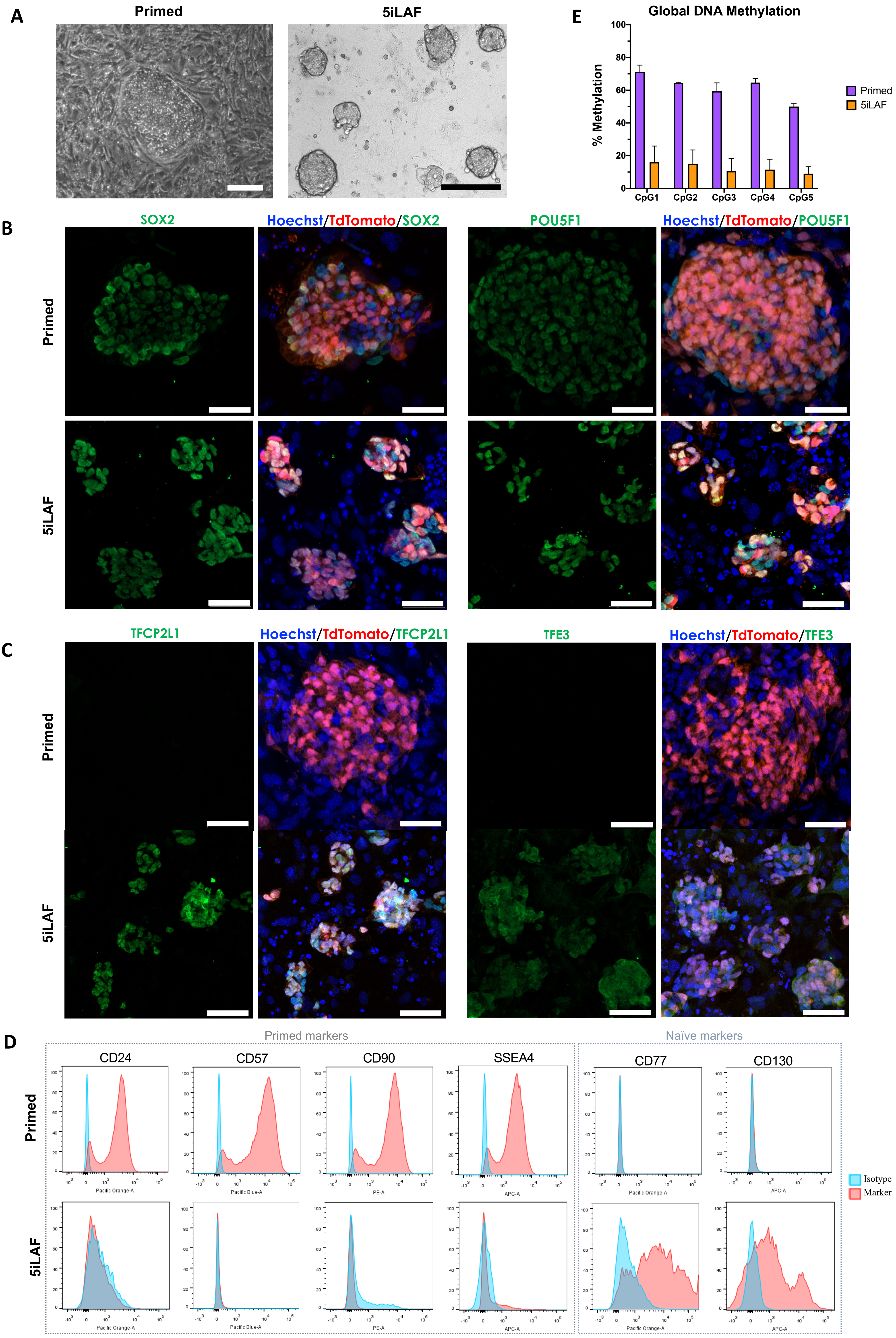

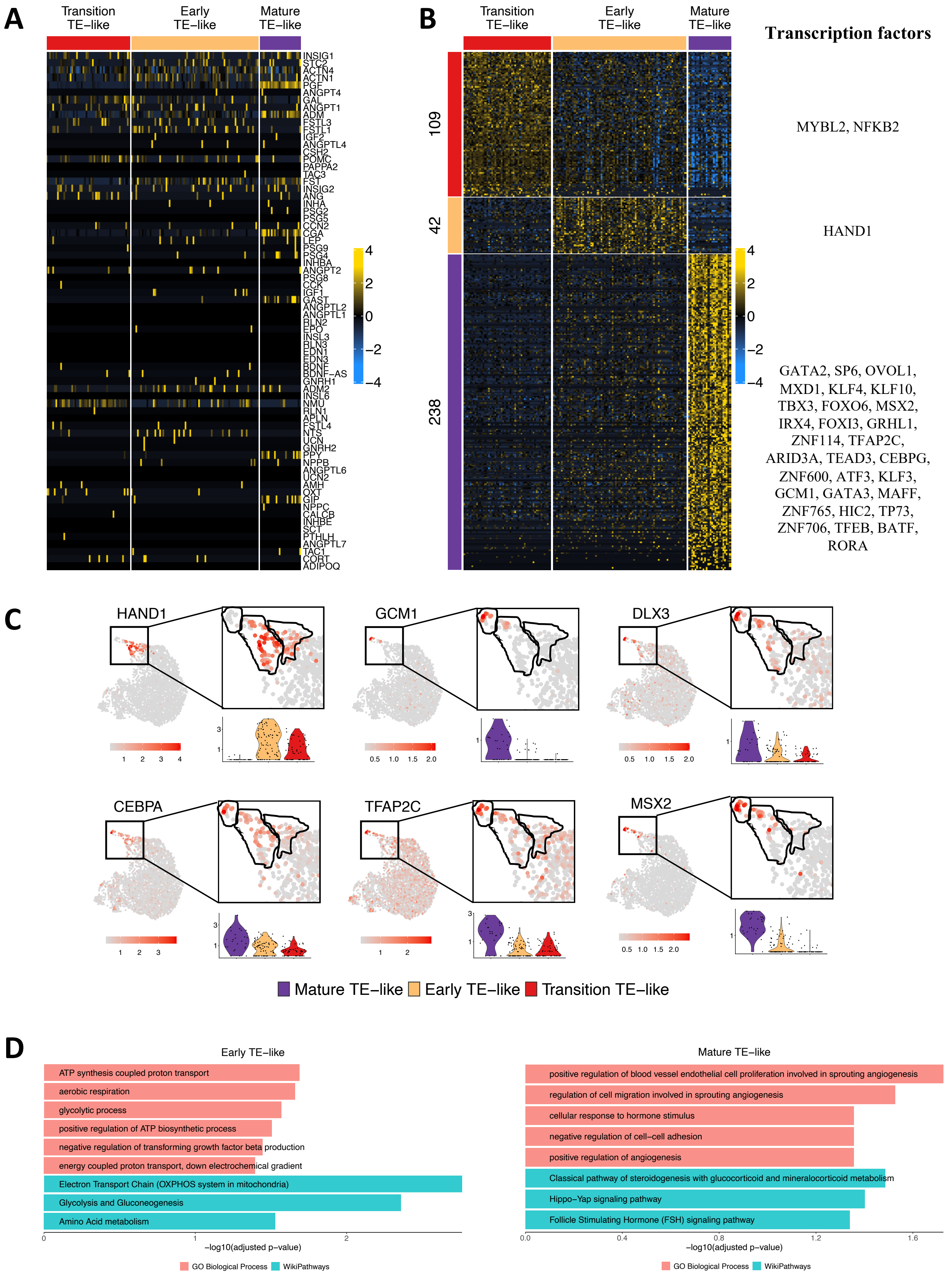
